# Fate mapping analysis reveals a novel dermal migratory Langerhans-like cell population

**DOI:** 10.1101/2020.12.10.419903

**Authors:** Jianpeng Sheng, Chen Qi, Dong Yu Wen, Wu Xiaoting, Johannes Mayer, Wang Lin, Bai Xueli, Liang Tingbo, Sung Yang Ho, Wilson Wen Bin Goh, Franca Ronchese, Christiane Ruedl

## Abstract

Dendritic cells residing in the skin represent a large family of antigen presenting cells, ranging from long-lived Langerhans cells (LC) in the epidermis to various distinct classical dendritic cell subsets in the dermis. Through genetic fate mapping analysis and single cell RNA sequencing we have identified a novel separate population of LC-independent CD207^+^CD326^+^ LC^like^ cells in the dermis that homed at a slow rate to the LNs. These LC^like^ cells were long-lived and radioresistant but, unlike LCs, they were gradually replenished by bone-marrow-derived precursors under steady state. LC^like^ cells together with cDC1s were the main migratory CD207^+^CD326^+^ cell fractions present in the LN and not, as currently assumed, LCs, which were barely detectable, if at all. These findings bring new insights into the dynamism of cutaneous dendritic cells and opens novel avenues in the development of treatments to cure inflammatory skin disorders.

## Introduction

In 1868, Paul Langerhans described a novel dendritic-shaped, non-pigmentary cell population in the epidermis (Langerhans, 1868). These so-called Langerhans cells (LCs) were first classified as cellular members of the nervous system, due to their morphological similarity with neurons. It was not until the 1980s when it became clear that this peculiar epidermal cell fraction with its potent antigen presentation activity belonged to the dendritic cell (DC) family (Romani and Schuler, 1989; Schuler and Steinman, 1985). Despite the fact that LCs share many features with DCs, they are generally considered as epidermal tissue-resident macrophages, mainly due to their dependence on CSF1, their embryonic origin and local self-maintenance (Wynn et al., 2013), although a conventional “macrophage signature” (e.g. CD16/32, CD64 and MerTK expression) is missing (Gautier et al., 2012).

LCs can sense invading pathogens and initiate an intrinsic maturation process that drives their migration out of the epidermis (Romani et al., 2001). As such, LCs have been regarded as a prototype antigen presenting cell (APC) (Nagao et al., 2009) that can, after antigen capture, migrate to the draining lymph nodes (LNs) to initiate an immune response by priming naïve LN-resident T cells (Romani et al., 2003). Antigen presentation can, however, occur in skin-draining LNs independently of LCs (Henri et al., 2010). In fact, the skin hosts several other distinct dermal DC subpopulations (Henri et al., 2010; Kissenpfennig et al., 2005), the presence of which complicates the analysis of the cellular contribution to skin immune responses, such as contact hypersensitivity (CHS). Consequently, the paradigm of “who is doing what” (i.e. epidermal LCs versus dermal DC counterparts) is still controversial (Bennett et al., 2005; Bobr et al., 2010; Bursch et al., 2007; Clausen and Stoitzner, 2015; Kaplan et al., 2005; Noordegraaf et al., 2010; West and Bennett, 2017).

Here we demonstrate that under steady-state conditions LCs most likely do not exit the skin, or if so, in very low numbers. Through a combined use of genetic fate mapping and novel inducible LC ablating mouse models, we show that the originally described LN LC fraction, is actually an independent LC^like^ cell population that originates from the dermis, not from the epidermis. These LC^like^ cells are ontogenitically different from LCs and are replaced over time by BM-derived cells with slow kinetics before trafficking to the LN.

## Results

### LC^like^ cells are found in dermis and LNs

The skin and the skin-draining LNs contain several distinct dendritic cell subpopulations. To delineate migratory LCs and dermal DCs, we profiled DC subsets in the epidermis, dermis and skin-draining LNs. In the epidermis, we confirmed that CD326^+^CD207^+^ LCs are predominantly found within the CD11b^hi^F4/80^hi^ fraction (Nagao et al., 2009; Valladeau et al., 2000) (Fig. 1A). In the dermis, we found a fraction of CD11b^hi^F4/80^hi^ cells that co-expressed CD326 and CD207 (Fig. 1B, upper panel). These cells could be immigrated LCs, although we can’t exclude a contamination from the epidermis during the isolation procedure. As expected, the remaining dermal CD11b^hi^F4/80^hi^ cells were CD326^-^CD207^-^ tissue-resident macrophages (Sheng et al., 2015; Tamoutounour et al., 2013). Dermal DCs were localized in the F4/80^int^ and CD11c^hi^MHCII^+^ DC fraction, which we could separate into three subpopulations based on CD103 and CD11b expression: CD103^+^CD11b^-^ (defined as cDC1), CD103^-^CD11b^low^ and CD103^-^CD11b^hi^ (defined as CD11b^hi^). CD103^+^CD11b^-^ DCs but not CD103^-^CD11b^hi^ DCs co-expressed CD326 and CD207. We could also divide the CD103^-^CD11b^low^ subpopulation into CD326^-^CD207^-^ (defined as Triple negative-TN) and CD326^+^CD207^+^ (defined as LC^like^) fractions (Fig. 1B, right panel). To track the corresponding migratory DCs in the skin-draining LNs, we first gated on CD11c^int-hi^MHCII^hi^ cells, which represent the migratory DC fraction (Sheng et al., 2017). Similar to our findings in the dermis, CD11b and CD103 labelling separated the migratory DCs into CD103^+^CD11b^-^ (cDC1), CD103^-^CD11b^hi^ (CD11b^hi^) and CD103^-^CD11b^low^ cells (Fig.1C). The CD103^-^CD11b^low^ cells could be further separated in two fractions: CD326^-^CD207^-^ (TN), and CD326^+^CD207^+^ (LC^like^) subpopulations (Fig. 1C). Notably, we did not detect the *bona fide* epidermal and dermal LCs showing the original F4/80^hi^CD11b^hi^ phenotype in the LN (Fig. 1C, right, lower panel). In agreement with previous work (Henri et al, 2010), the CD11b^hi^ DC fraction represented the largest DC subpopulation in the dermis, whereas in the LN all four DC subpopulations (CD11b^hi^, cDC1, TN and LC^like^) were almost equally represented (Fig. 1D). Because we detected no phenotypic F4/80^hi^ LCs in the LNs, we hypothesized that the cutaneous DCs en route to the LN were not derived from epidermal LCs, but rather from distinct dermal CD11b^hi^, cDC1, TN and F4/80^low^ LC^like^ DC populations. This analysis can’t exclude, however, the possibility that the migrating LCs might change their phenotype.

**Fig. 1:**
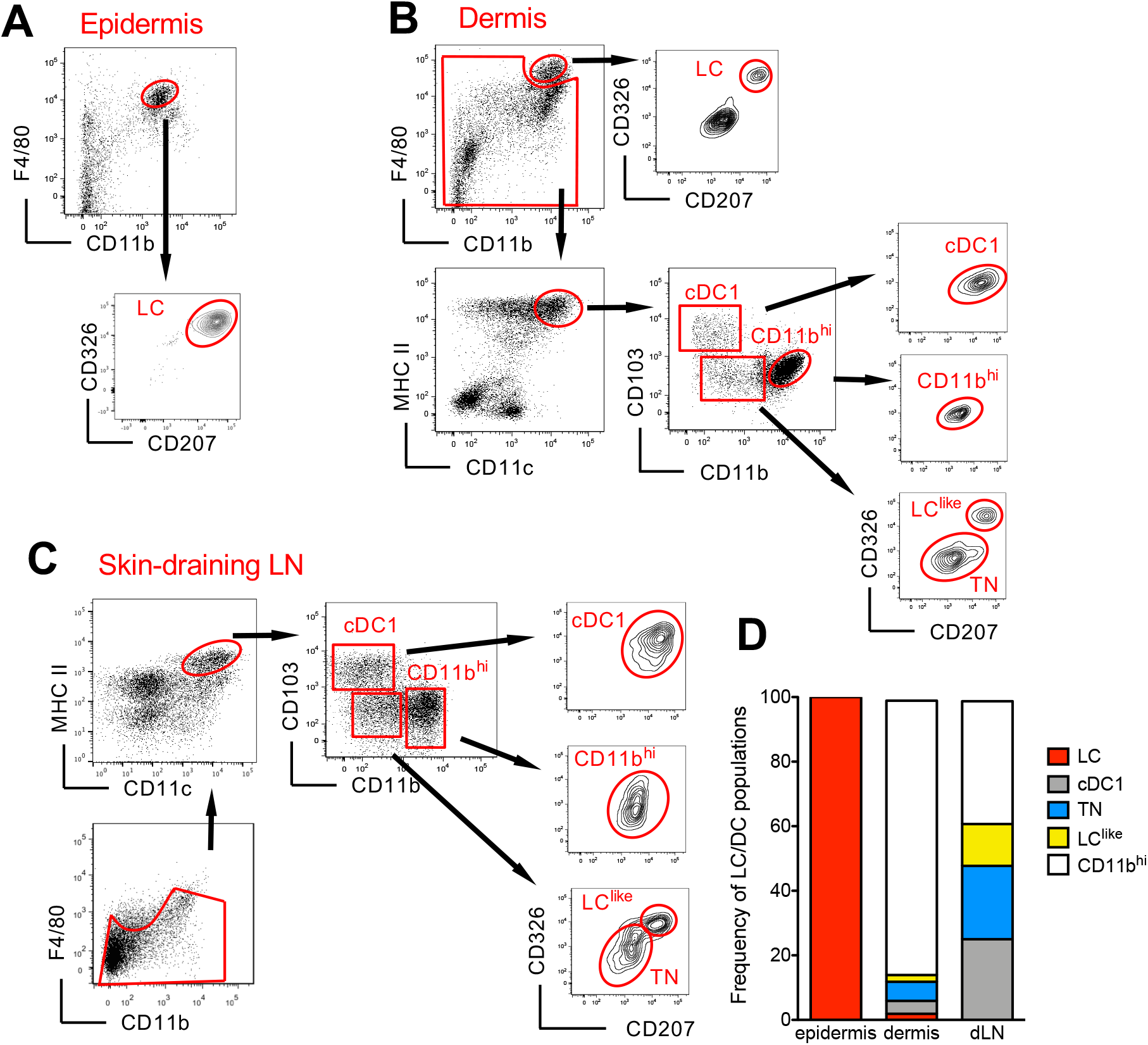
Characterization of cutaneous LC and DC subpopulations. **A**,Representative flow cytometry dot plots for LC characterization in the epidermis. Cells from epidermis were first gated for FSC, SSC and CD45 (not shown). Then, CD45+ cells were analyzed for CD11b and F4/80 expression. The CD11b^hi^F4/80^hi^ cell fraction was further analysed for CD207 and CD326 expression to identify classical *bona fide* LCs. **B,**Representative flow cytometry dot plots for dermal LC and DC subpopulations. Isolated dermis cells were first gated for FSC, SSC and CD45 (not shown). CD45+ cells were then analyzed for CD11b and F4/80 expression. The CD11b^hi^ F4/80^hi^ fraction contained classical CD207^+^CD326^+^ LCs. The remaining cells were gated for CD11c^+^MHC II^+^ DCs and separated into three subsets based on CD103 and CD11b expression: CD103^+^CD11b^-^ cells (labelled cDC1), CD103^-^CD11b^hi^ DCs (labelled CD11b^hi^), and CD11b^low/neg^. CD207 and CD326 expression was detectable on cDC1 but not CD11b^hi^ DCs, whereas CD11b^low^ cells were further separated into CD207^-^CD326^-^ (labelled TN) and CD207^+^CD326^+^ (labelled LC^like^). **C,**Representative flow cytometry dot plots for cutaneous DC subpopulations in skin-draining LNs. LN cells were first gated for FSC and SSC before F4/80 and CD11b staining. The cell fraction excluding F4/80^hi^/CD11b^hi^ cells was separated by CD11c and MHCII. CD11c^hi^MHCII^hi^ migratory DCs were gated and analysed for CD103, CD11b, CD207 and CD326 expression. Four subsets were detected: CD103^+^CD11b^-^CD207^+^CD326^+^ (cDC1), CD103^-^CD11b^low^CD207^-^CD326^-^ (TN), CD103^-^CD11b^low^CD207^+^CD326^+^ (LC^like^) and CD103^-^CD11b^hi^CD207^-^CD326^-/+^(CD11b^hi^). **D,**, Frequency of each DC subpopulation (LC, cDC1, LC^like^, TN and CD11b^hi^) present in epidermis, dermis and cutaneous LN, respectively.

### Single cell RNAseq confirms the presence of two independent LC and LC^like^ cell populations in the dermis

Since LC and LC^like^ cells co-exist together in the dermis, we aimed to investigate their relationship and respective gene signature by scRNAseq analysis. Unsupervised clustering and uniform manifold approximation and projection (UMAP) projections were performed on 9605 enriched cells isolated from the dermis of ears obtained from 5 mice. The origin of distinct CD45^+^ and CD45^-^ dermal cell subpopulations are visualized in a color-coded UMAP plot (Fig. 2A). Nine different cell clusters could be broadly identified by unsupervised clustering and classified as follows: (1) LC, (2) LC^like^, (3) Mast cells/neutrophils, (4), DC/monocytes, (5) macrophages, (6) lymphocytes 1, (7) lymphocytes 2, (8) mesenchymal cells and (9) epithelial cells. Conventional DCs, monocyte and other myeloid related signature genes, such as *zbtb46* (DCs), *xcr1* and *clec9A* (cDC1), *siglec H* (plasmacytoid DC), *ly6c* and *ccr2* (monocytes), *gata2* and *FcεrI* (mast cells) and *ly6g* (neutrophils) are mainly detectable in the DC/mono and mast cell/neutrophil clusters (3-4) and are mainly absent or weakly expressed in the LC/LC^like^ clusters (1-2) (Fig. 2B, C and Fig. S1). *cd207* and *cd326* expressing cells are detected in LC (1), LC^like^ (2) as well as in DC/monocyte cluster (4) which confirms the presence of three distinct CD207^+^CD326^+^ dermal, subpopulations observed by flow cytometry (Fig. 1B). *Cd207 cd326* expressing cells detected in the cluster 4 are co-expressing *clec9A, xcr1, irf8* hence they represent the cDC1s (Fig. 2B and Fig. S1). *Cd207 cd326* expressing cells in clusters 1 (LC) and 2 (LC^like^) share many of previously reported LC signature genes (e.g. *cd11c, f4/80, cd74, mafb, spi1, csf1r, tgfbr1)* (Fig. 2C and Fig. S1), but several other genes are differentially expressed in LC^like^ cells (e.g. *tgfbr2, sylt3, col27a1, fernt2, spry2)* or in LC cells (e.g. *cd209a, agpat4, birc3, dusp16, gdpd3, ly75 and ppfibp2),* respectively (Fig. 2C, D). In summary, the unsupervised clustering of single cells obtained from dermis suggests that LC and LC^like^ cells are two independent cell fractions and distinct from CD207^+^EpCAM^+^ cDC1s as already shown in conventional flow cytometry analysis (Fig 1B).

**Fig. 2:**
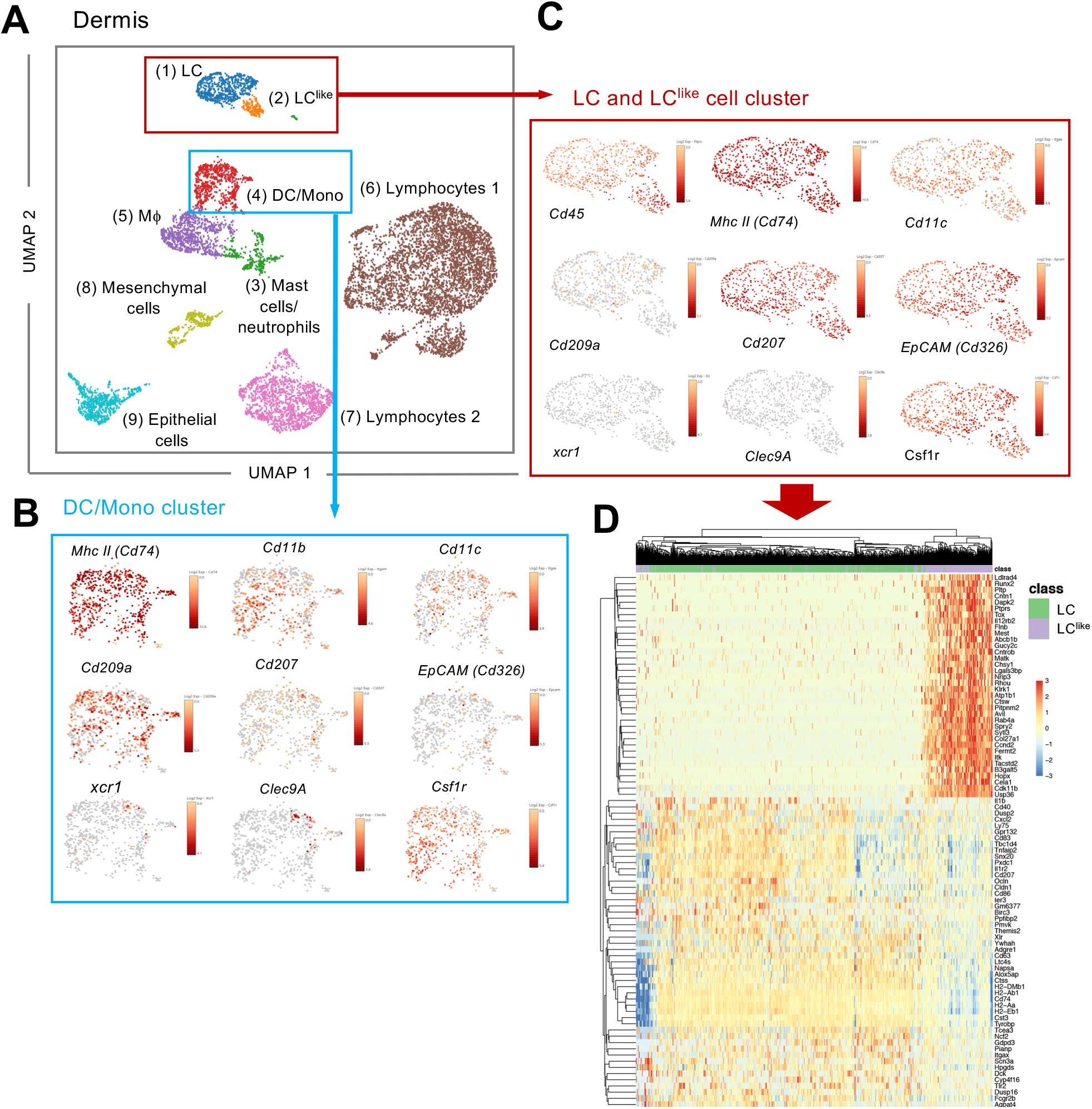
Single cell RNAseq analysis reveals LC and LC^like^ cells as two distinct cell populations in the dermis. 9605 cells pooled from the dermis collected from 6 mice which passed QC were imported for Seurat analysis. **A,**UMAP plot is revealing the existence of 9, distinct cell clusters (1) LC [blue], (2) LC^like^ [orange], (3) Mast cells/neutrophils [green], (4) DC/monocytes [red], (5) macrophages [purple], (6) lymphocytes 1 [brown], (7) lymphocytes 2 [pink], (8) mesenchymal cells [light green] and (9) epithelial cells [light blue]. **B, C**, UMAP maps showing the expression of various LC signature genes in DC/mono **(B)**and LC/LC^like^ clusters **(C)**. **D,**heat-map of single-cell gene expression data based on the top differential expressed genes discriminating LC/LC^like^ clusters. Cells (LC in green; LC^like^ in purple) are shown in rows and genes in columns.

### Early yolk sac precursors contribute to the development of LC but not LC^like^ cells

Fate mapping experiments have shown that epidermal LCs derived partially from primitive yolk sac progenitors (Hoeffel et al., 2012, Sheng et al., 2015), therefore the developmental origin of LCs is distinct from conventional DCs and resembled more microglia. To study in detail a possible yolk sac origin of distinct cutaneous LC and DC subpopulations, a single injection of TAM was given to E7.5 pregnant *Kit*^MerCreMer/R26^ mice (Fig. 3A). Three months later the epidermis, dermis and brain (microglia as positive control) were collected and isolated cells were then analysed for YFP expression. As previously reported, microglia, the prototype yolk sac derived macrophage, were strongly labeled (~40%) (Fig. 3B, E). However, about 12 % of epidermal LCs were YFP labelled, confirming their partial yolk sac origin (Fig. 3C, E). In comparison, the dermal LC counterparts showed a similar labeling profile (~10%), whereas the remaining dermal DC subpopulations (LC^like^, cDC1, CD11b^hi^ and TN) showed a significantly lower 5% YFP signal, very likely, attributed to small spill over of labeling in the HSCs (Fig. 3D, E). Therefore, YS only contributed to LCs but not to LC^like^ cells.

**Fig. 3:**
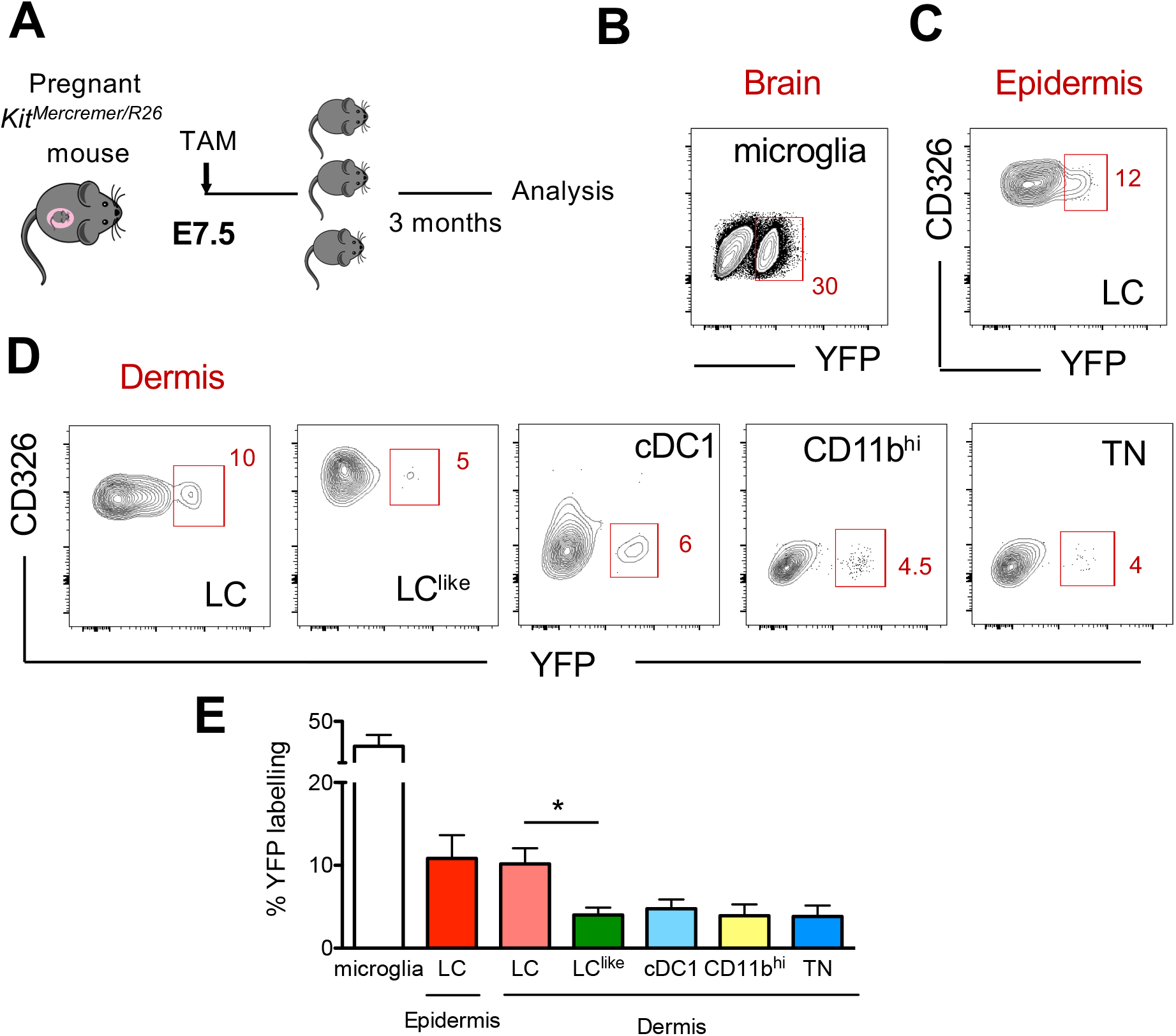
Distinct embryonic origin between LC and LC^like^ cells. **A,**Single pulse of TAM at E7.5 was given to label Kit^MercreMerR26^ embryos and the percentages of labeled brain microglia (positive control), epidermal LCs and dermal LC/DC subpopulations were measured at 3 months of age. **B-D,**Flow cytometry analysis of YFP labelling of microglia **(B)**, and each LC and DC subpopulation in the epidermis **(C)**and dermis **(D)**in *Kit*^MerCreMer/R26^ fate mapping mice. Representative contour plots are shown. **E,**The mean percentage of YFP^+^ cells of brain microglia, epidermal LC and dermal DC subpopulations (LC, cDC1, LC^like^, CD11b^hi^ and TN cells). The error bars represent the SEM (n= 4 samples of 2-3 pooled mice). Data from two independent experiments. * P <0.05; two-way ANOVA followed by Bonferroni test. For clarity non-significant values are not shown.

### LC^like^ DCs exhibit dual origins

LCs are the only cell type from the DC family that originate from self-renewing radio-resistant embryonic precursors (Merad et al., 2002); other DC subpopulations are short-lived and constantly replenished by BM-progenitors ^23^. To delineate the radioresistant properties of the newly identified LC^like^ cells, we generated bone marrow (BM) chimeric mice by transplanting congenic CD45.1^+^ mouse BM cells into irradiated CD45.2^+^ recipients (Fig. 4A). We then analysed the CD45.1^+^/ CD45.2^+^ ratio in different skin-related DC subpopulations 1 or 4 months after reconstitution.

**Fig. 4:**
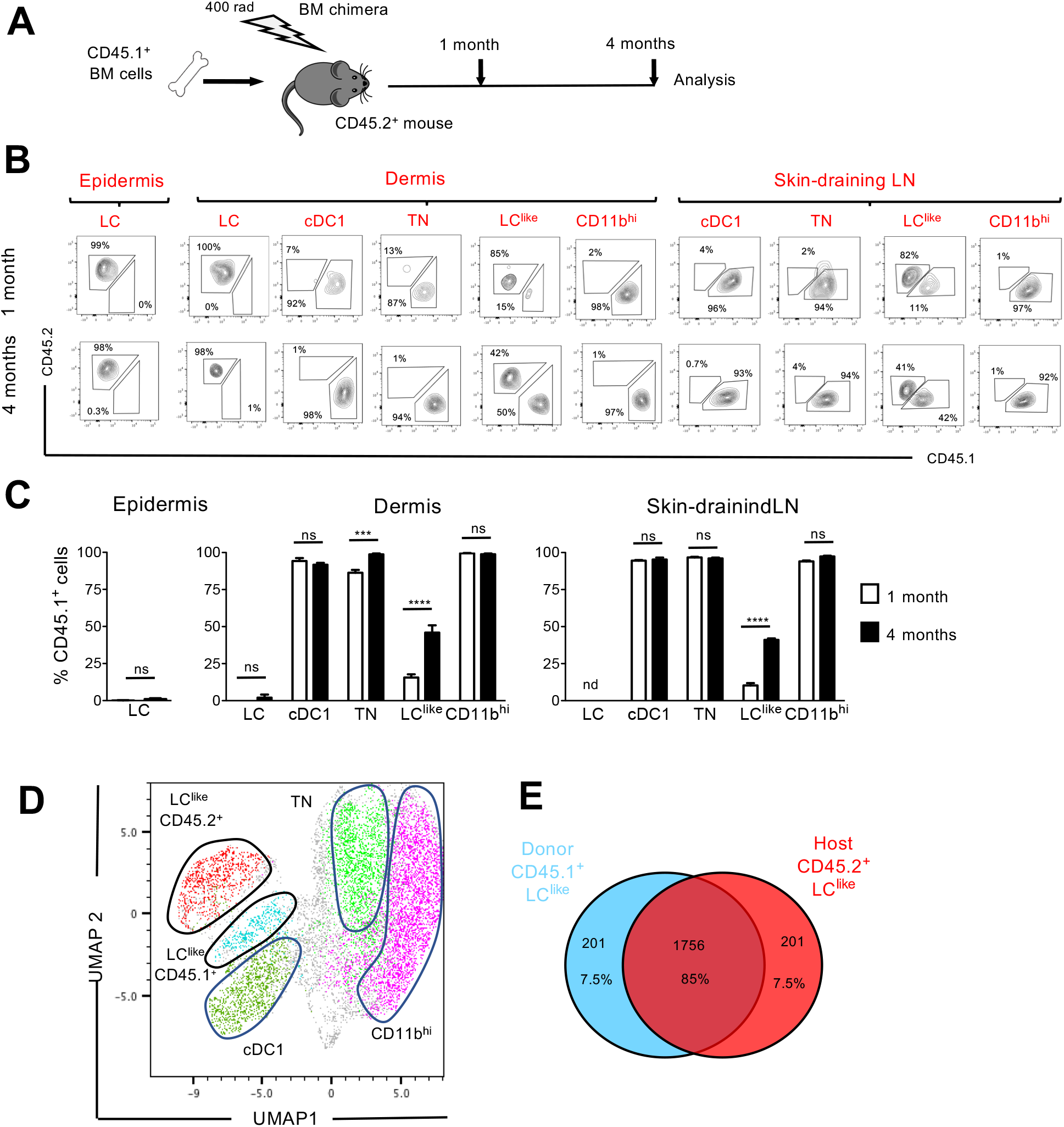
LC^like^ cells display a dual origin with a similar transcriptomic signature. **A,**Generation of BM chimeras: CD45.1^+^ WT BM cells (10^6^) were transferred into lethally irradiated CD45.2^+^ recipient mice. The epidermis, dermis and draining LNs obtained from the reconstituted chimeras were analysed 1 and 4 months later by flow cytometry. **B,**Flow cytometry analysis of donor (CD45.1^+^) and host (CD45.2^+^) chimerism in different epidermal, dermal and skin-draining LN LC and DC subpopulations, 1 and 4 months after reconstitution. LC, cDC1, TN, LC^like^ and CD11b^hi^ subsets were gated and analysed for CD45.1 (x-axis) and CD45.2 (y-axis) expression. **C,**The percentage of CD45.1 donor cells, detected in the epidermis, dermis and skin-draining LNs of chimeras, 1 or 4 months after reconstitution. Data are represented as mean ± SEM; *n* = 6 single mice; *** P< 0.001; **** P< 0.0001; ns, non-significant; unpaired Student’s t-test. **D,**UMAP analysis of distinct LN DC subpopulations obtained from chimeras 4 months after reconstitution, based on the expression of different markers (CD11c, MHCII, CD103, CD11b, EPCAM, CD207, CD45.1, CD45.2). **E,**Transcriptome analysis of LN CD45.1^+^ LC^like^ cells (n=3) and LN CD45.2^+^ LC^like^ (n=3) cells collected from 10 mice. The Venn diagram shows the percentage of overlapping genes expressed by CD45.1^+^ and CD45.2^+^ LC^like^ cells.

In the epidermis and dermis, LCs were mostly CD45.2^+^, and thus retained their host origins due to local self-renewal (Fig. 4B, C). By contrast, dermal cDC1, TN and CD11b^hi^ DCs exhibited a wholly CD45.1^+^ phenotype after just 1 month following reconstitution; this finding means that they are fully BM-derived. Only LC^like^ cells showed a mixed contribution from both CD45.2^+^ host and CD45.1^+^ donor cells. In fact, after 1 month following reconstitution, only a minority (~10%) of LC^like^ cells were replenished by CD45.1^+^ cells; this percentage increased to ~50% by 4 months after reconstitution (Fig. 4B, C).

In skin-draining LNs, we found that cDC1, TN and CD11b^hi^ cells were mostly derived from donor CD45.1^+^ BM cells, excluding their origins from the radio-resistant LC population. Comparable to its dermal counterpart, only the LC^like^ cell fraction was split into donor CD45.1^+^ and host CD45.2^+^ cells, respectively (Fig. 4B, C). In addition, the contribution of CD45.1^+^ donor cells increased over time, from ~10% after 1 month to ~45% after 4 months. This unique temporal replacement suggests a dual origin for LC^like^ cells, distinguishing this DC fraction from both conventional long-lived radio-resistant self-renewing LCs and short-lived BM-derived DCs.

To allow high resolution and unbiased data-driven dissection of skin DC subpopulations in the reconstituted chimeric mice, we performed a Uniform Manifold Approximation and Projection (UMAP) analysis of flow cytometry data. Both CD45.1^+^ and CD45.2^+^ LC^like^ cells were clearly visible and clustered separately but in close proximity (Fig. 4D). Using this dimensional reduction algorithm, we detected that CD11c^+^MHCII^hi^ dermal dendritic cell subpopulations could be grouped into five separate clusters: cDC1, TN, CD11b^hi^ and two LC^like^ cell clusters (BM-derived CD45.1^+^ and resident CD45.2^+^). To investigate the molecular relationship between the resident LC^like^ cell population and the BM-derived LC^like^ cells, we performed RNA-sequencing on LN LC^like^ cells isolated from chimeric mice (CD45.1^+^ donor BM cells into CD45.2^+^ recipient mice). Unsupervised hierarchical clustering (Euclidean Distance, Complete linkage) and Principle Component Analysis (PCA) analysis revealed that both CD45.1^+^ and CD45.2^+^ LC^like^ cells clustered closely together (not shown), with ~85 % of their gene expression overlapping (Fig. 4E). The high level of similarity between resident and BM-derived LC^like^ fractions indicates that the microenvironment and not the cellular origin, seems to determine the LC^like^ cell identity.

### LC^like^ cells display slow turnover kinetics

BM chimeras require full body irradiation, which can damage the local skin micro-environment and attract BM-derived newcomers. This irradiation could, therefore, complicate the analysis of skin resident cell homeostatic turnover kinetics. To circumvent this issue, we performed a fate-mapping study under steady state conditions using *Kit*^MerCreMer/R26^ fate mapping mice. These mice allow for the turnover rates of cell populations derived from BM precursors to be estimated (Sheng et al., 2015). We performed our analyses at different time points (1, 4 and 8 months) after TAM injection to ensure a sufficiently long time-frame to monitor populations that turn over slowly (Fig. 5A).

**Fig. 5:**
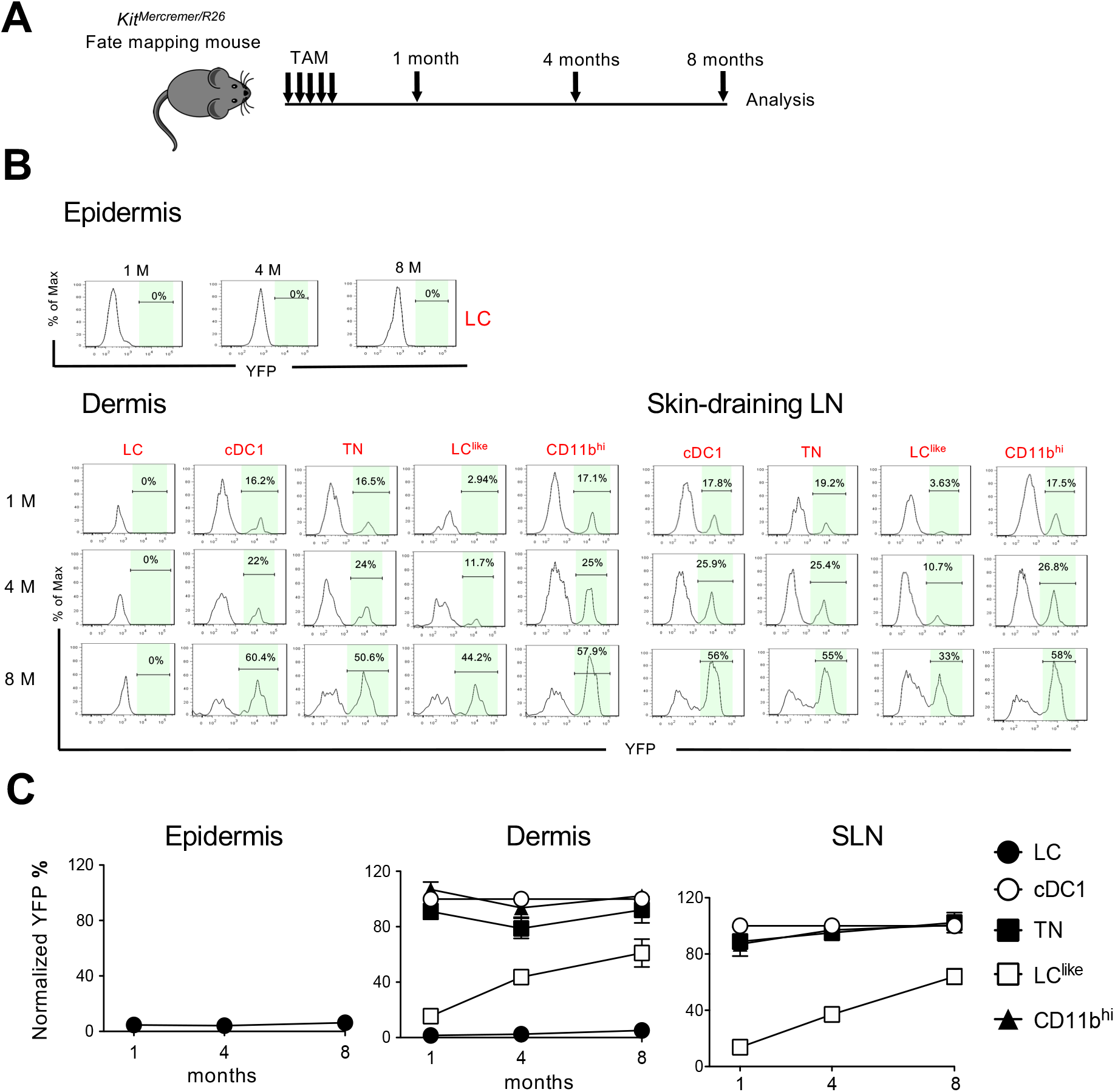
Slow turnover kinetics for dermal and LN LC^like^ cells. **A,**Kit^MerCreMer/R26^ mice aged 6-weeks old mice were injected with tamoxifen five times and groups of six animals were sacrificed 1, 4 and 8 months later for fate mapping analysis. **B,**Flow cytometry analysis of YFP labelling of each LC and DC subpopulation in the epidermis (upper left), dermis (lower left) and skin-draining LNs (lower right) in *Kit*^MerCreMer/R26^ fate mapping mice. Representative histograms are shown. **C**, The mean percentage of YFP^+^ cells after normalization to cDC1. Epidermis (left), dermis (middle) and skin-draining LNs (right) were analysed. Data are represented as mean ± SEM; *n* = 6-9 single mice. **** P <0.0001; two-way ANOVA followed by Bonferroni test.

In the epidermis, LCs showed minimal YFP labelling over the entire 8-month chase period; this finding was expected as these cells are not replaced by BM-derived cells. (Fig. 5B and C, left panel). Similarly in the dermis, CD11b^hi^F4/80^hi^CD326^+^CD207^+^ cells showed minimal labelling from 1 to 8 months (Fig. 5B and C, middle panel). We propose that this fraction most likely represents immigrant LCs in the dermis. cDC1, TN and CD11b^hi^ DCs, however, were fully labelled with YFP after just 1 month and the labelling was maintained for the remaining 8 months. This finding is consistent with the fast turnover rate identified for these three DC subsets. By contrast, LC^like^ cells gradually accumulated the label from 10 to 60% over the 8-month chase period, supporting that dermis-resident LC^like^ DCs are replaced slowly by BM progenitors. In the skin-draining LNs, all DC subsets behaved similarly to their dermal counterparts (Fig. 5B and C, right panel). Briefly, cDC1, TN and CD11b^hi^ DCs showed a fast turnover by reaching plateau level of labelling after 1 month while LC^like^ cells demonstrated a slow turnover rate over the 8-month chase period.

### LC^like^ cells are not derived from classical LCs

To interrogate the relationship between LC and LC^like^ cells, we exploited a novel DC-SIGN-DTR transgenic mouse strain (Fig. S2A). which allowed us to deplete epidermal and dermal LCs without affecting the LC^like^ cell pool and other immune cells types (Fig. 6 and Fig. S2B). In fact, we detected DC-SIGN (or CD209a) by qPCR in murine LCs and in CD11b^hi^ DCs but not in cDC1, TN or LC^like^ cells (Fig. S2C), a result which was corroborated by the scRNA analysis (Fig. 2). We established short and long depletion protocols (Fig. 6A) to capture even potentially very slowly migrating “LC-derivatives” (Bursch et al., 2007). In the DT-treated DC-SIGN DTR mice, LCs were efficiently depleted in both the epidermis and dermis by the short-term and long-term depletion protocols (Fig. 6A–C, Fig. S2D). We also found that cells in the CD11b^hi^ cell fraction were affected by the DT treatment; this was particularly evident during the short-term depletion protocol, in which the cell numbers were reduced by ~80 % (Fig. 6B, C). Importantly, cDC1, TN and LC^like^ cell numbers were unaffected and thus were comparable between DT-injected WT and DC-SIGN mouse strains. These results strongly support the independency of LC^like^ cells from classical *bona fide* epidermal LCs.

**Fig. 6:**
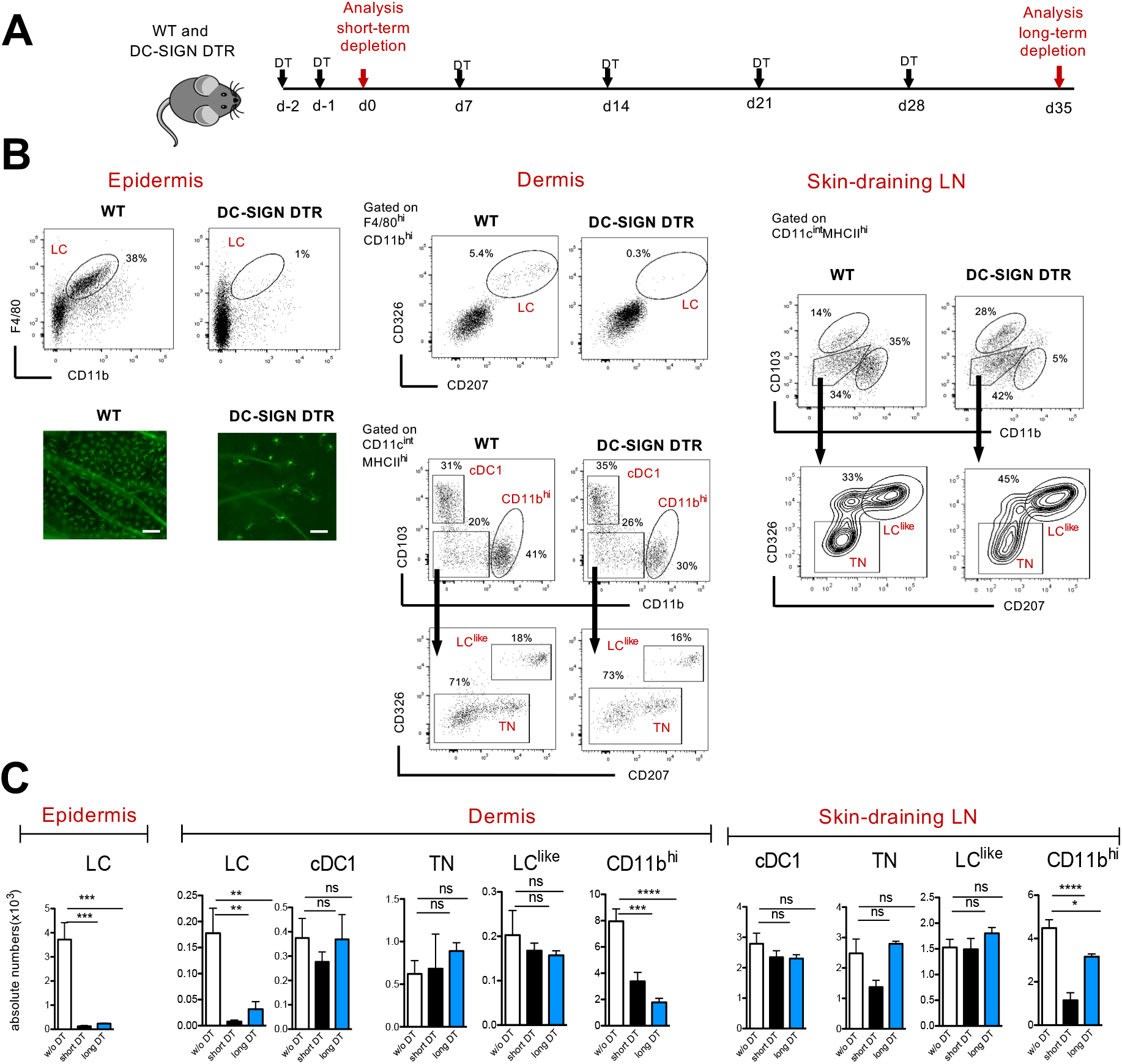
Classical LC, but not LC^like^ cells are ablated in vivo in DC-SIGN DTR mice. **A,**The short-term and long-term depletion protocol in DC-SIGN-DTR mice. **B,**Representative flow cytometry dot plots of single-cell suspensions from the epidermis (left), dermis (middle) and skin-draining LNs (right) obtained from DT-injected WT and DC-SIGN-DTR mice. All mice were injected (i.p.) with 10 ng/g DT on days −2 and −1 and analysed on day 0. The gating strategy shown in Figure 1 was followed. Epidermal sheets obtained from WT and DC-SIGN mice were stained for MHC class II (green fluorescence) and analysed by immunofluorescence microscopy (lower left panel). **C,**The absolute numbers of each gated myeloid cell subset (LC, cDC1, TN, LC^like^ and CD11b^hi^ cells), obtained from the epidermis, dermis and skin-draining LNs of DT-injected WT and DC-SIGN DTR mice. Data are represented as mean ± SEM; *n* = 8 single mice. **** P, <0.0001; *** P <0.001; * P <0.05; ns, non-significant; one-way ANOVA followed by Bonferroni test.

To further confirm that LC^like^ cells represent a distinct cell lineage from LCs, we crossed DC-SIGN DTR mice with a *Kit*^Mercremer/R26^ fate mapping mouse, which would enable us to trace BM-derived cells in absence of LC. We treated these mice (DC-SIGN DTR-*Kit*^Mercremer/R26^) with TAM and then injected them with DT for 5 weeks to maintain long-term LC depletion (Fig. 7A). Although epidermal LCs were absent over the whole period, the YFP labelling profiles of skin-derived LN DC subsets, including the LC^like^ fraction, were comparable between DT-injected DC-SIGN DTR^+^-*Kit*^Mercremer/R26^ and DC-SIGN DTR^neg^-*Kit*^Mercremer/R26^mice (Fig. 7B, C). These data show that in absence of LC the replenishment of LC^like^ cells by BM-derived cells is not affected.

**Fig. 7:**
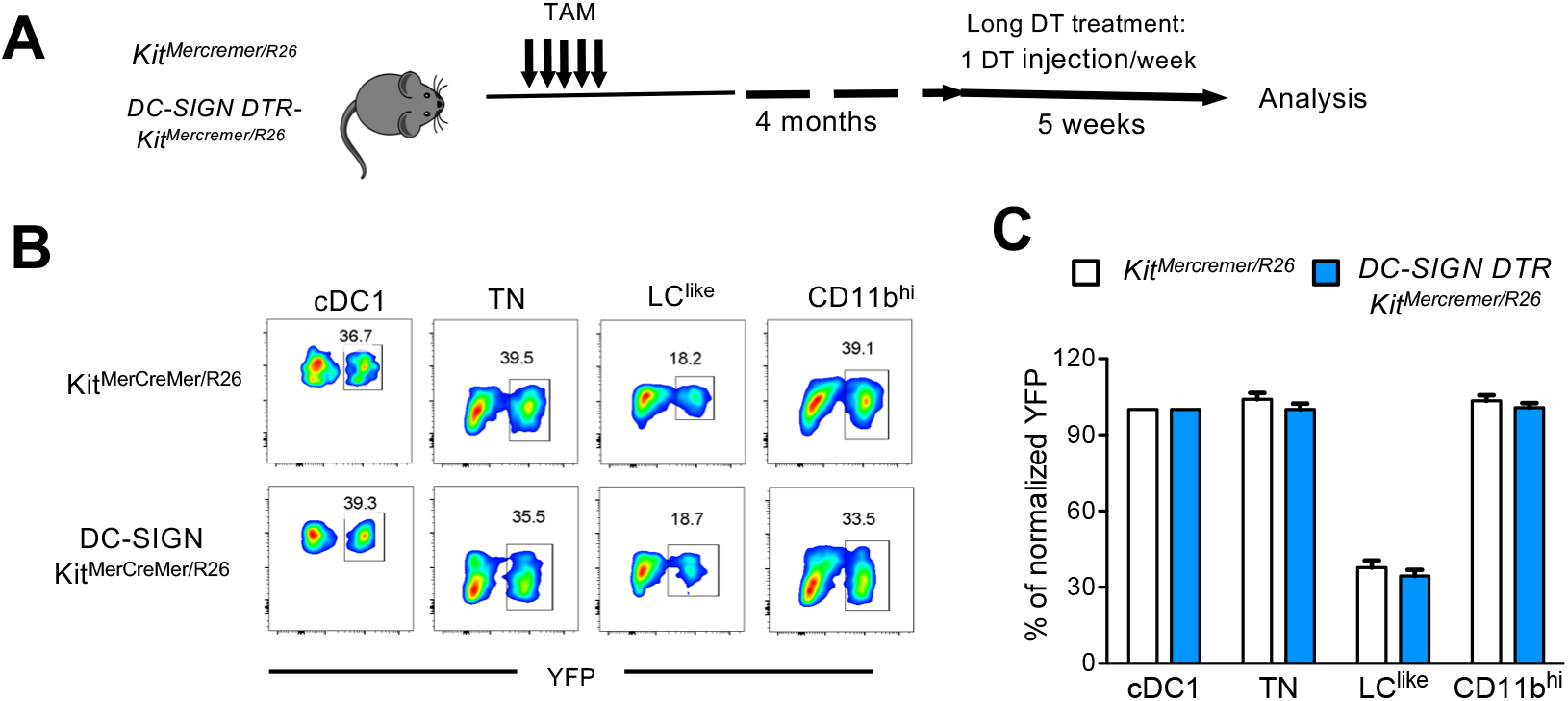
Fate mapping analysis in DC-SIGN DTR-Kit^MerCreMer/R26^ mice. **A,**Mice aged 6-weeks old were orally gavaged with TAM. After 4 months, DT was injected i.p. weekly for 5 weeks to ensure long-term LC depletion. **B,**Representative contour plots showing the YFP labelling of distinct LN DC subpopulations in DT-treated Kit^MerCreMer/R26^ and DC-SIGN DTR-Kit^MerCreMer/R26^ mice. **C,**The percentage of normalized YFP labelling detected in DC subpopulations (LC, cDC1, TN, LC^like^ and CD11b^hi^ cells) of the skin-draining LNs. Normalization was performed as described in Fig. 5; Data are represented as mean ± SEM; *n* = 12 single mice.

## Discussion

Epidermal Langerhans cells (LCs) are the only antigen presenting cells localized in the epidermis. These cells were recently re-defined as “macrophages in dendritic cell clothing” due to their unique ontogeny, and self-renewing and radioresistant characteristics (Doebel et al., 2017). By contrast, there are multiple DC and macrophage subpopulations that reside in the dermis (Tamoutounour et al., 2013). Although these dermal DCs share some common markers with LCs [such as langerin (CD207) and EpCaM (CD326)], they constitute a distinct cell lineage on the basis of their developmental origins and cytokine requirements (Bursch et al., 2007; Ginhoux et al., 2007; Poulin et al., 2007). Three CD207^+^ DC subpopulations have been described in the skin-draining LN: two subpopulations are skin-derived and one subpopulation originates from the BM (Bursch et al., 2007; Douillard et al., 2005; Henri et al., 2001; Romani et al., 2010). Due to this diverse skin-resident DC network, it became evident that not only LCs, but other skin-derived DCs might be involved either in tolerance or immune response induction in draining LNs.

Although it is commonly believed that the journey of a LC starts from the epidermis and ends in the skin-draining LN after a transit through the dermis in steady state, we found that it is in-fact their look-alike counterparts, LC^like^ cells, that migrate to the draining LNs. Our new insight was gained by re-analysing established mouse strains (*Kit*^MerCreMer/R26^ mice) and exploiting newly generated transgenic mouse strains (DC-SIGN-DTR mice and DC-SIGN-DTR-Kit^MerCreMer/R26^ fate mapping mice), which allowed us to visualize, with increasing resolution, the *in vivo* dynamics of skin-resident DCs under steady state. We first characterized and redefined different DC/LC subsets in the dermis by flow cytometry and scRNAseq analysis, which delineated classical F4/80^hi^ LC and four different DC subsets, namely cDC1, TN DCs, CD11b^hi^ DCs and an unappreciated CD11b^low^F4/80^low^ LC^like^ cell fraction. With the exception of classical F4/80^hi^ LCs, we found all of these cells in the migratory CD11c^int^MHCII^hi^ DC fraction of the skin-draining LN. This finding suggests that the majority of migratory CD326^+^CD207^+^ DCs are CD103^+^ cDC1 and CD103^-^ LC^like^ cells and not classical CD11b^hi^F4/80^hi^ LCs which are hardly seen in the LN if not at all.

Corroborating evidence for differential migratory behaviours among different skin dendritic cells was provided by real-time intravital two-photon microscopy. Under steady-state conditions, due to the structural integrity of the basement membrane, epidermal LCs are sessile with static and almost immobile dendrites. In contrast, dermal DC subpopulations are actively crawling through the dermal interstitial space at high velocity even in absence of inflammation suggesting that continuous migration to LN is a steady-state property of dermal DCs and not epidermal LCs (Ng et al., 2008; Shklovskaya et al., 2010).

Our analysis of dermal DCs is in full agreement with Henri et al. who similarly to us disentangled the DC family in the dermis in five subpopulations: two subsets lacking the expression of CD207 (CD207^-^CD11b^-^ [TN] and CD207^-^CD11b^+^ [CD11bhi]) and three expressing CD207 (CD11b^int^CD207^++^ mLCs, CD11b^low/-^CD207^+^CD103^+^ [cDC1] and CD11b^low^CD207^+^CD103^-^ [LC^like^]) (Henri et al., 2010).

Similarly, cutaneous LNs, were distinguished in five analogous subpopulations including mLCs, which were defined for their characteristics in radioresistance and not for the expression of classical LC markers (CD11b^hi^ and F4/80^hi^). Henri et al. speculated that LN LCs downregulated CD11b and F4/80 expression (Henri et al., 2010) and therefore these markers lost their discriminatory power to segregate distinct CD207^+^ cells in the LN.

To circumvent the “complication” of the potential shift in phenotype, we adopted an alternative approach based on genetic fate-mapping analyses which allowed to trace cell lineages between distinct LC and DC subpopulations avoiding lethal irradiation and generation of chimeric mice. First, our E7.5 embryo “labelling strategy” demonstrated that only LCs are partially yolk sac derived (Sheng et al, 2015), but not the other migratory DCs subpopulations, inclusive LC^like^ cells. Second, our detailed analyses of the fate mapping kinetics revealed that the radioresistant and radiosensitive CD11b^low^CD207^+^CD103^-^ subpopulations described by Henri et a. represented instead a truly homogeneous radioresistant LC^like^ subpopulation, which is gradually replaced over time by BM-derived progenitors. Furthermore, we corroborated their “LC-independency”, since long-term absence of LCs did not affect the numbers of LC^like^ cells in our DC-SIGN DTR mouse, model. Accordingly, our analysis delineated only four, and not five, LN migratory DC subpopulations, excluding LCs.

The phenotypes, transcription profiles and cytokine requirements of dermal cDC1, TN and CD11b^hi^ DCs have been extensively described [reviewed in (Clausen and Stoitzner, 2015)]; however, there has been comparatively less attention given to the LC^like^ subpopulation. Unsupervised clustering of scRNA-seq trascriptome data of dermal cells indicated that LC and LC^like^ cells, although sharing some common myeloid cell markers, are two independent cell fractions and clearly distinct from macrophages and the other skin DC subpopulations. However, unlike LCs, which are BM-independent, radio-resistant and self-renewing (Ghigo et al., 2013; Hoeffel et al., 2012), LC^like^ cells represent a radio-resistant population that is progressively replaced postnatally by BM-derived precursors. Similar to resident macrophages in tissues, such as skin, gut, kidney and heart, LC^like^ cells have a dual origin involving both embryonic (but not yolk-sack like LCs) and adult haematopoiesis (Molawi et al., 2014; Sheng et al., 2015; Soncin et al., 2018). Unlike other skin DC subpopulations, which are short-lived and exhibit a high turnover rate, we show that fetal-derived LC^like^ cells are long-lived and are replaced very slowly by BM-derived cells. These fetal-derived and BM-derived LC^like^ cells co-exist together in adult tissue, and although derived from different origins, they show high similarity. This finding suggests that it is the local tissue microenvironment and not the cellular origin that shapes their final identity. The existence of a LC-independent radio-resistant dermal DC fraction was previously observed in other study that described the presence of an *in situ* proliferating, radio-resistant dermal DC subpopulation not only in the murine but also in human dermis (Bogunovic et al., 2006). It is likely that these cells are the LC^like^ cells described here.

Although all dermal DCs migrate into LNs in a CCR7-dependent fashion (Forster et al., 1999), LC^like^ cells seem to migrate at slower rate than other DCs under steady state conditions. Similar slow trafficking dynamics was originally attributed to LCs (Bursch et al., 2007; Ruedl et al., 2000) but we now strongly believe that these previously reported slow migratory cells are in fact LC^like^ cells.

To further rule out the possibility that epidermal F4/80^hi^CD11b^hi^ LC downregulate CD11b and F4/80 and turn into F4/80^low^CD11b^low^ LC^like^ cells in the dermis, we exploited a novel DC-SIGN DTR transgenic mouse strain where LCs, but not LC^like^ cells, could be ablated. Even long-term depletion (6 weeks) of epidermal and dermal LCs had no effect on the numbers of LC^like^ cells in the dermis and LNs while maintaining their LC^like^ YFP-labeling profile in the absence of epidermal LCs in DC-SIGN DTR-*Kit*^Mercremer/R26^ mice.

In summary, our genetic fate-mapping approach, used to delineate the complex skin DC network, does not support the established paradigm of LCs as being the main “prototype” migrating APCs to draining LNs under homeostatic conditions (Wilson and Villadangos, 2004). We propose, rather, that LCs at steady state, similar to other tissue-resident macrophages, are sessile and act locally in the skin, whereas dermal LC^like^ cells assume many of the functions previously attributed to LCs. The identification of this novel migratory dermal LC^like^ subpopulation opens new avenues and approaches in the development of treatments to cure diseases such as contact allergic dermatitis and other inflammatory skin disorders like psoriasis.

## Material and Methods

### Mice

C57BL/6J and B6.SJL-*Ptprc^a^ Pepc^b^*/BoyJ (B6 CD45.1) were obtained from The Jackson Laboratory (USA). *Kit*^MerCreMer/Rosa26-LSL-eYFP^ (called *Kit*^MerCreMer/R26^) mice were generated as previously described ((Piva et al., 2012; Sheng et al., 2015). Kit^MerCreMer/R26^ mice were backcrossed with DC-SIGN-DTR mice to obtain DC-SIGN-DTR-Kit^MerCreMer/R26^ mice.

DC-SIGN DTR mice were generated as follows: the IRES-DTR fusion gene was inserted into the 3’-UTR region of the DC-SIGN gene locus on BAC RP24-306K4; the gene targeting vector was then retrieved from the modified BAC (Fig. S2A). The gene targeting vector was linearized and electroporated into Balb/C embryonic stem (ES) cells and correctly recombined ES colonies were selected by PCR. Gene targeted ES cells were injected into C57BL/6 blastocysts and transferred into the oviduct of a pseudo-pregnant mother. F0 male chimera mice were mated with F1 Balb/C females to obtain F1 Balb/C DC-SIGN DTR mice; these mice were then back crossed to C57BL/6 for 12 generations to generate C57BL/6 DC-SIGN DTR mouse.

All mice were bred and maintained in the specific pathogen-free animal facility of the Nanyang Technological University (Singapore). All studies involving mice in Singapore were carried out in strict accordance with the recommendations of the National Advisory Committee for Laboratory Animal Research and all protocols were approved by the Institutional Animal Care and Use Committee of the Nanyang Technological University. For animal work performed in New Zealand, experimental protocols were approved by the Victoria University of Wellington Animal Ethics Committee and performed in accordance with institutional guidelines.

### Tamoxifen-inducible fate-mapping mouse models

Kit^MerCreMer/R26^ and DC-SIGN-DTR-Kit^MerCreMer/R26^ fate-mapping mice were used to monitor the turnover rates of distinct skin-related DC subpopulation subsets. Each mouse was, administered 4 mg tamoxifen (TAM) (Sigma-Aldrich, St. Louis, MO, USA) for five consecutive days by oral gavage for adult labelling, as previously described (Sheng et al., 2015). Pregnant mice (E7.5) were injected once with 16 mg TAM for embryo labelling.

### Diphteria toxin (DT) injection

DC-SIGN-DTR^pos^ and DC-SIGN-DTR^neg^ mice were injected intraperitoneally (i.p.) with 20 ng/g DT (Sigma-Aldrich) to deplete DC-SIGN-expressing cells. Two different DT injection protocols were used (Fig. 6A). For the short term depletion protocol, mice were injected i.p. at day −2 and −1 before collections of tissues. For the long-term protocol, DT was injected once a week over 5 weeks prior tissue collection.

### Generation of BM chimeras

Chimeric mice were generated by irradiating recipient C57BL/6 or DC-SIGN-DTR mice (CD45.2^+^) with two doses of 550 Gy, 4 h apart. Then, 10^6^ B6.Ly5.1 (CD45.1^+^) BM cells were injected intravenously (i.v), 24 h after treatment. The mice were allowed to recover from 1 to 4 months before analysis.

### Isolation of epidermal, dermal and LN cells

Mouse ears were cut and separated into dorsal and ventral halves using fine forceps. Both the dorsal and ventral halves (with the epidermis side facing upwards) were incubated for 1 h at 37°C in 1 ml IMDM (Thermo-Fisher Scientific, Waltham, MA, USA) medium containing 1U Dispase II (Thermo-Fisher Scientific). The epidermis and dermis were separated using fine forceps, cut into small pieces and digested for another 1 h at 37°C in 1 mg/ml Collagenase D (Roche, Basel, Switzerland). To obtain single-cell suspensions, the digested tissue was passed through a 40 μm cell strainer. To process skin-draining LNs, the dissected LNs were minced and incubated in 1 mg/ml Collagenase D for 60 min at 37°C.

### Antibodies

The following antibodies were used: anti-mouse CD45 (30-F11), anti-mouse CD11b (M1/70), anti-mouse F4/80 (BM8), anti-mouse Ly6c (HK1.4), anti-mouse CD11c (N418), anti-mouse I-A/I-E (M5/114.15.2), anti-mouse CD103 (2E7), anti-mouse CD326 (G8.8), anti-mouse CD207 (4C7), anti-mouse CD45.1 (A20), anti-mouse CD45.2 (104). They were purchased all from Biolegend (San Diego, CA, USA). Anti-mouse CD45 microbeads from Milteny (Bergisch Gladbach, Germany).

### Flow cytometry analysis of skin-related DC subpopulations

Single-cell epidermal, dermal or LN tissue suspensions were pre-incubated with 10 μg/ml anti-Fc receptor antibody (2.4G2) on ice for 20 min. Then, the suspensions were further incubated with fluorochrome-labeled antibodies at 4°C for 20 min, before being washed and re-suspended in PBS/2% FCS for analysis on a five-laser flow cytometer (LSR Fortessa^TM^; BD Bioscience, San Jose, CA, USA). The data were analysed with FlowJo software (TreeStar, Ashland, OR, USA) and UMAP analysis was performed using the FlowJo UMAP plugin.

### Single cell RNAseq analysis

Immune cells were enriched using anti-mouse CD45 microbeads from dermal single cell suspension. Briefly, enriched CD45^+^ dermal cells were loaded into Chromium microfluidic chips with v3 chemistry and barcoded with a 10× Chromium Controller (10X Genomics, Pleasanton, CA, USA). RNA from the barcoded cells was subsequently reverse-transcribed and sequencing libraries constructed with reagents from a Chromium Single Cell v3 reagent kit (10X Genomics) according to the manufacturer’s instructions. Library sequencing was performed at Novogene Co., Ltd (Tianjin Novogen Technology Co., Tianjin, China) with Illumina HiSeq 2000 according to the manufacturer’s instructions (Illumina, San Diego, USA).

### Single cell data analysis

FastQC was used to perform basic statistics on the quality of the raw reads. Raw reads were demultiplexed and mapped to the reference genome by 10X Genomics Cell Ranger pipeline using default parameters. All downstream single-cell analyses were performed using Cell Ranger and Seurat unless mentioned specifically. In brief, for each gene and each cell barcode (filtered by Cell Ranger), unique molecule identifiers were counted to construct digital expression matrices. Secondary filtration for Seurat analysis: a gene with expression in more than 3 cells was considered as expressed and each cell was required to have at least 200 expressed genes.

### RNA-sequencing analysis

All mouse RNAs were analyzed using an Agilent Bioanalyser (Agilent, Santa Clara, CA, USA). The RNA Integrity Number (RIN) ranged from 3.4-9.3, with a median of 8.2. cDNA libraries were prepared from a range of 18, 24.2, 68 and 100 ng total RNA starting material using the Ovation Universal RNA-seq system. The length distribution of the cDNA libraries was monitored using a DNA High Sensitivity Reagent Kit on an Agilent Bioanalyser. All 11 samples were subjected to an indexed paired-end sequencing run of 2×100bp on an Illumina Novaseq 6000 system (Illumina, San Diego, CA, USA).

The paired-end reads were trimmed with trim_galore1 (option: −q 20 –stringency 5 –paired). The trimmed paired-end reads were mapped to the Mouse GRCm38/mm10 reference genome using the STAR2 (version 2.6.0a) alignment tool with multi-sample 2-pass mapping. Mapped reads were summarized to the gene level using featureCounts3 in the subread4 software package (version 1.4.6-p5) and with gene annotation from GENCODE release M19. DESeq25 was used to analyze differentially expressed genes (DEGs), and significant genes were identified with Benjamini-Hochberg adjusted P-values <0.05. DESeq2 analysis was carried out in R version 3.5.2.

For functional analysis, hierarchical clustering based on Euclidean distance and complete linkage, was performed using the R “pheatmap” package. Principal Components Analysis, (PCA) was performed using the R “prcomp” package. The first two Principal Components (PC) were analysed on a multi-dimensional scatterplot that was created using the R “scatterplot 3D” function.

### Preparation and staining of epidermal sheets

DC-SIGN-DTR^neg^ and DC-SIGN-DTR^+^ mice were treated for 2 days with DT. Ears were collected and split into dorsal and ventral halves and subsequently incubated with 3.8% ammonium thiocyanate (Sigma-Aldrich) in PBS for 20 min at 37°C. Epidermal and dermal sheets were separated and fixed in ice-cold acetone for 15 min. Then, the epidermal sheets were pre-incubated with 10 μg/ml anti-Fc receptor antibody (2.4G2) on ice for 20 min and subsequently stained with FITC-labelled anti MHC class II antibody for a further 30 min on ice for LC visualization.

### Statistics

The data represent the means ± SEM or SD, as indicated in the Figure Legends. GraphPad Prism software was used to display the data and for statistical analysis. Statistical tests were selected based on the appropriate assumptions with respect to data distribution and variance characteristics. All statistical tests are fully described in detail in the Figure Legends. Samples were analyzed by two-tailed Student’s *t*-test to determine statistical differences between two groups. A two-way ANOVA with Bonferroni-post-test was used to determine the differences between more than two groups. A P value <0.05 was considered to be statistically significant. The number of animals used per group is indicated in the Figure Legends as ‘‘n.’’

## Data and Code Availability

All RNA-sequencing data have been deposited in the Gene Expression Omnibus public database under accession number GSE139877. Single cell RNAseq have been deposited into NCBI SRA database with BioProject ID: PRJNA625270.

Original flow cytometry data are deposited in the NTU Open Access Data Repository DR-NTU. All other data are available from the authors upon reasonable request.

## Author Contributions

Conceptualization: J.S and C.R.; Methodology: J.S., Q.C., D.Y., W.X., J.M., F.R; Formal Analysis: J.S and C.R.; Bioinformatic analysis: Y.H.S. and W.B.W.G.; scRNAseq analysis: W.L., B.X. and L.T. Writing: J.S. and C.R. Visualization: C.R. Supervision: C.R. Funding Acquisition: J.S. and C.R.

## Acknowledgements

The authors would like to thank Monika Tetlak for mouse management, Su I-hsin and Lim Jun Feng Thomas for technical advice and Insight Editing London for proofreading the manuscript prior to submission. This work was supported by Ministry of Education Tier1 grant awarded to C.R. and by the National Key R&D Program of China (Grant 2019YFA0803000) assigned to J.S․. Work at the MIMR was supported by an Independent Research Organisation grant awarded to the Malaghan Institute of Medical Research from the Health Research Council of New Zealand

---

**Fig. S1:**
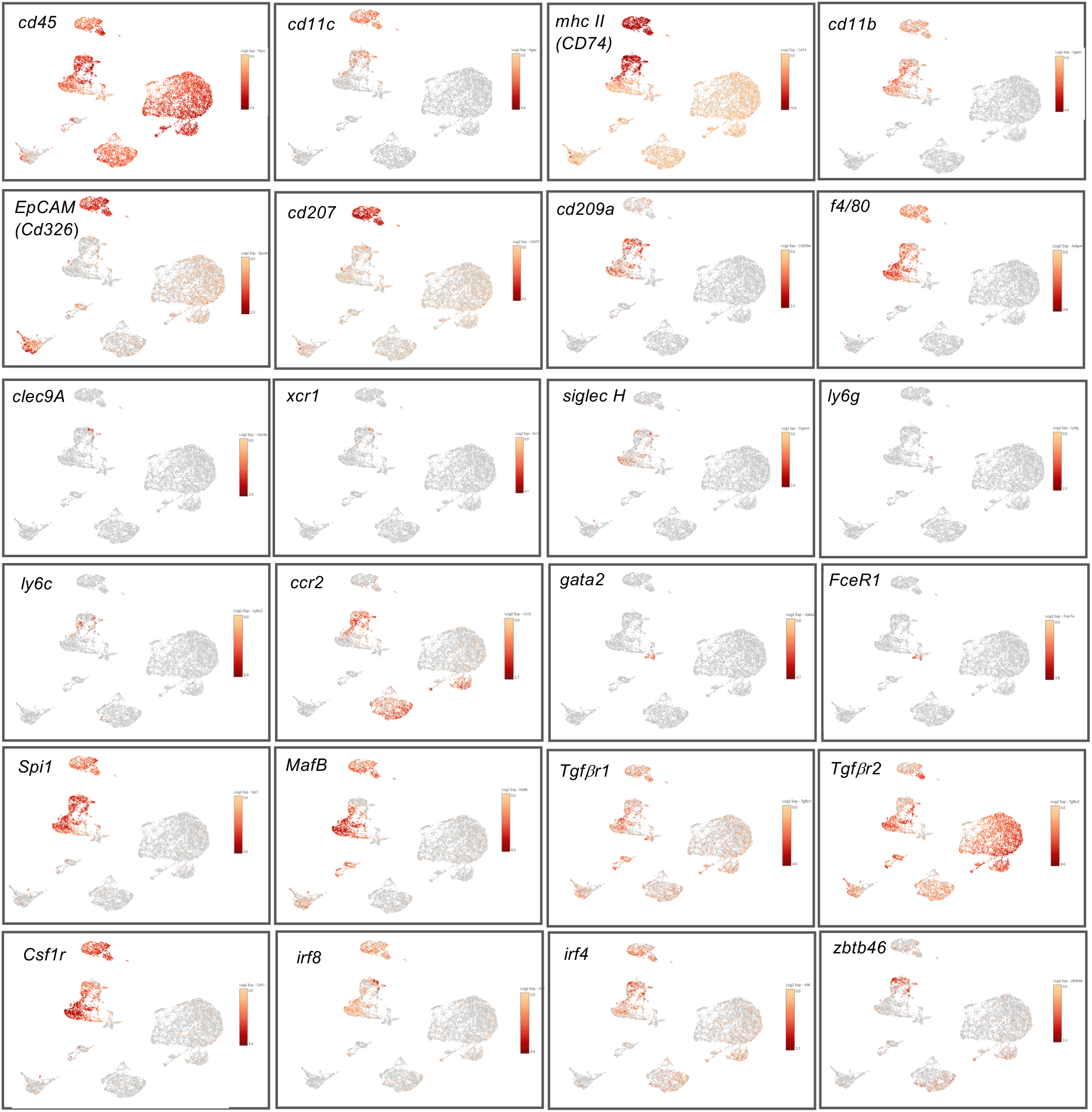
Analysis of scRNAseq data from murine dermis. UMAP plots showing the expression of indicated prototypical myeloid genes.

**Fig. S2.**
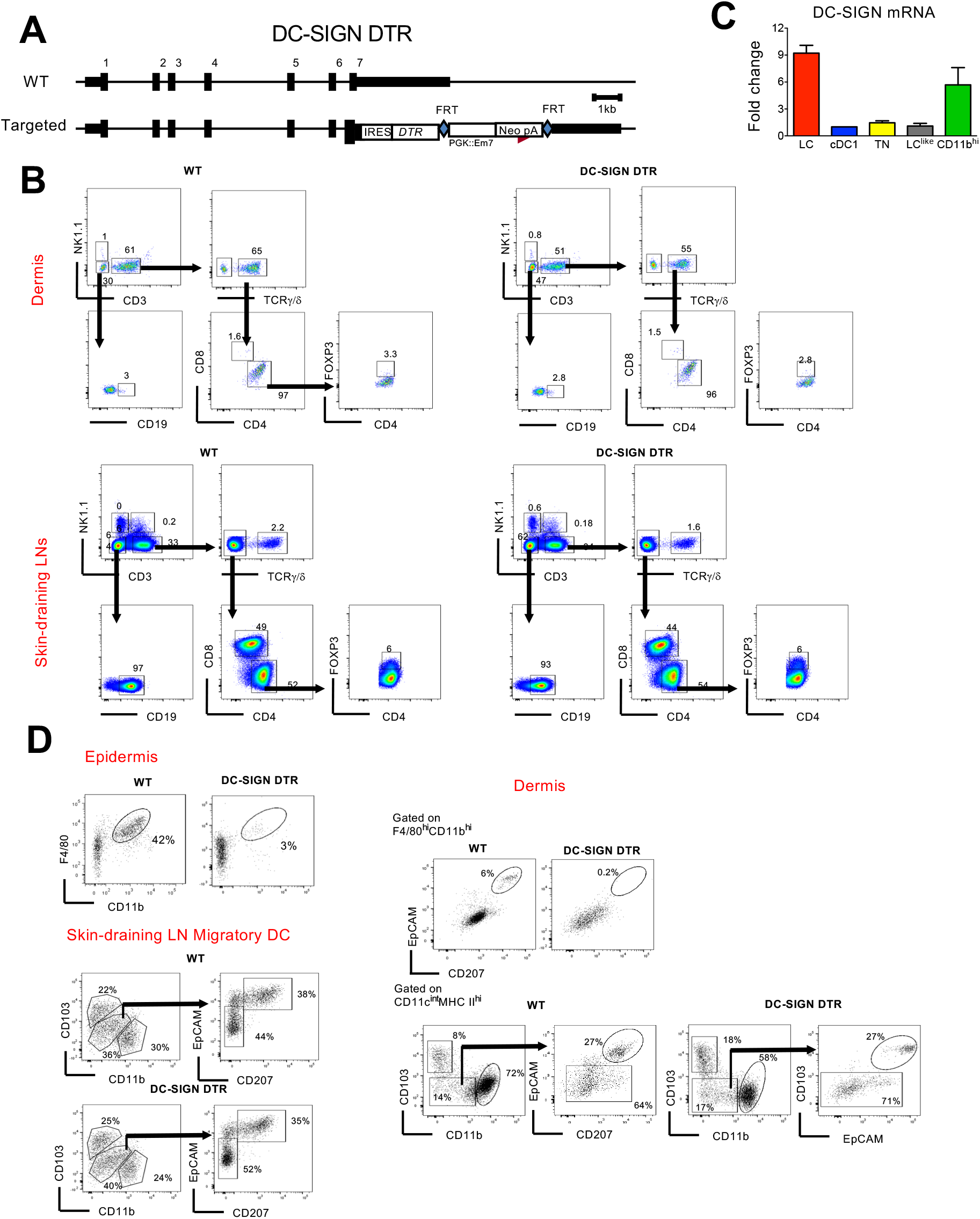
(A) The experimental strategy to generate DC-SIGN-DTR mice. (B) Quantitative PCR analysis of the mRNA expression in epidermal LC and distinct LN DC subpopulations. (C) Lymphoid cell populations in DT-treated WT and DC-SIGN-DTR mice. Upper panel: dermis; lower panel: skin-draining LNs. (D) Representative flow cytometry dot plots of single-cell suspensions from the epidermis (top), dermis (middle) and skin-draining LN (bottom) obtained from DT-injected WT and DC-SIGN-DTR mice following the long-term depletion protocol shown in Fig. 6A.

